# High-resolution reconstruction of cell-type specific transcriptional regulatory processes from bulk sequencing samples

**DOI:** 10.1101/2025.04.02.646189

**Authors:** Li Yao, Sagar R. Shah, Abdullah Ozer, Junke Zhang, Xiuqi Pan, Tianyu Xia, Vrushali D. Fangal, Alden King-Yung Leung, Meihan Wei, John T. Lis, Haiyuan Yu

**Affiliations:** Department of Computational Biology, Cornell University, Ithaca, NY 14853; Weill Institute for Cell and Molecular Biology, Cornell University, Ithaca, NY 14853; Department of Molecular Biology and Genetics, Cornell University, Ithaca, NY 14853

## Abstract

Biological systems exhibit remarkable heterogeneity, characterized by intricate interplay among diverse cell types. Resolving the regulatory processes of specific cell types is crucial for delineating developmental mechanisms and disease etiologies. While single-cell sequencing methods such as scRNA-seq and scATAC-seq have revolutionized our understanding of individual cellular functions, adapting bulk genome-wide assays to achieve single-cell resolution of other genomic features remains a significant technical challenge. Here, we introduce Deep-learning-based DEconvolution of Tissue profiles with Accurate Interpretation of Locus-specific Signals (DeepDETAILS), a novel quasi-supervised framework to reconstruct cell-type-specific genomic signals with base-pair precision. DeepDETAILS’ core innovation lies in its ability to perform cross-modality deconvolution using scATAC-seq reference libraries for other bulk datasets, benefiting from the affordability and availability of scATAC-seq data. DeepDETAILS enables high-resolution mapping of genomic signals across diverse cell types, with great versatility for various omics datasets, including nascent transcript sequencing (such as PRO-cap and PRO-seq) and ChIP-seq for chromatin modifications. Our results demonstrate that DeepDETAILS significantly outperformed traditional statistical deconvolution methods. Using DeepDETAILS, we developed a comprehensive compendium of high-resolution nascent transcription and histone modification signals across 39 diverse human tissues and 86 distinct cell types. Furthermore, we applied our compendium to fine-map risk variants associated with Primary Sclerosing Cholangitis (PSC), a progressive cholestatic liver disorder, and revealed a potential etiology of the disease. Our tool and compendium provide invaluable insights into cellular complexity, opening new avenues for studying biological processes in various contexts.

## Introduction

Transcription is a tightly regulated process involving multiple rate-limiting steps, including the binding of transcription activators, chromatin remodeling and modification, enhancer-promoter interactions, preinitiation complex (PIC) formation, promoter escape, promoter-proximal pausing, and release of paused Pol II into productive elongation^1,2^. Understanding the regulatory mechanisms governing transcription is crucial for deciphering complex biological processes.

High-throughput sequencing assays, which capture snapshots of various regulatory aspects, have enhanced our understanding of these processes. For example, chromatin immunoprecipitation followed by sequencing (ChIP-seq) maps genome-wide distributions of transcription factor binding and histone modifications^3^. Chromatin conformation capture assays^4^, such as Hi-C, can catch enhancer-promoter interactions. Nascent transcript sequencing assays such as PRO-seq^5^, 3′ of CoPRO^6^, and mNET-seq^7^ can measure the dynamics of polymerase pausing and escape. The interplay between regulatory layers^8^ makes it possible to repurpose assays designed for profiling one process to capture the convoluted signal from other processes. For example, GRO/PRO-cap^5,9^ were originally designed for precise annotation of all active transcription start sites (TSSs) genome-wide, but the recent discovery regarding transcription at enhancer loci^10^ makes them highly sensitive assays for pinpointing active enhancers^11^. Collectively, these bulk sequencing technologies provide powerful tools to study the transcriptional regulatory processes that control cellular function and fate.

While powerful, bulk sequencing assays inherently aggregate signals across the entire cell populations, which can obscure cell-type-specific regulatory mechanisms when applied to tissue samples usually composed of diverse cell types that are available in limited quantities. In contrast, single-cell sequencing technologies such as single-cell/single-nucleus RNA sequencing (sc/snRNA-seq) and single-cell/single-nucleus assay for transposase-accessible chromatin sequencing (sc/snATAC-seq)^12,13^ offer a transformative approach by resolving cellular heterogeneity and enabling the characterization of regulatory processes at the individual cell level. In addition to measuring mRNA expression and chromatin accessibility, recent technological advances in this field also enabled the detection of more genomic features^14–18^.

However, adapting bulk sequencing assays to single-cell resolution in general faces significant experimental challenges, including the scarcity of input material and limitations in the efficiency of library preparations which can hinder certain assay functionalities^18^. Moreover, ease of use^15,16^ and cost^19^ present additional obstacles.

Alongside the development of single-cell-resolution assays, there is increasing interest in developing in silico deconvolution methods to reconstruct cell-type-specific profiles from sequencing data generated from bulk samples. Many current methods primarily rely on single-cell sequencing data from one assay type as the reference to deconvolute bulk sequencing data from the same modality. A widely explored approach involves developing statistical tools to estimate cell-type fractions^20,21^ and cell-type-specific gene expression from bulk RNA-seq samples using gene count matrices from scRNA-seq as references^22–26^. Due to the need for a panel of either bulk or reference samples, these tools were primarily used on consortium-level data such as TCGA^22,23^ and SFARI^25,27^. Recently, efforts have expanded to deconvolve other modalities, such as DNA methylation^28,29^, using references from the same modalities. However, obtaining single-cell sequencing reference libraries for many other omics modalities (e.g. nascent transcriptome) remains costly and challenging. Given the interconnected nature of regulatory layers, an alternative approach is to leverage single-cell references from more widely available modalities, such as sc/snATAC-seq, to deconvolve bulk data from assays that are not single-cell-ready (i.e. cross-modality deconvolution). However, a major challenge in cross-modality deconvolution is the lack of direct correspondence or alignment between different omics measurements, which can lead to discrepancies in data interpretation. For instance, a genomic region may exhibit similar accessibility across multiple cell types but only initiate transcription in one. This scenario is unique to cross-modality analyses, as intra-modality deconvolution does not encounter such complexities due to the inherent alignment within the same data type. As a result, we believe statistical methods will not be sufficient and we need to develop a new type of methods to both estimate the profiles of cell types and model the differences between the modalities.

In this study, we developed DeepDETAILS, a novel deep learning model enabling precise deconvolution of bulk omics profiles into cell-type-specific signals at base-pair resolution in a cross-modality manner. DeepDETAILS deconvolves bulk sequencing data—such as nascent transcriptome and histone modification profiles—using a sc/snATAC-seq library from the same type of tissue as reference. Our results demonstrate that DeepDETAILS significantly outperformed traditional statistical deconvolution methods. Using DeepDETAILS, we developed a comprehensive compendium of high-resolution nascent transcription and histone modification maps across 39 human tissues and 86 distinct cell types. Furthermore, we applied our compendium to fine map risk variants associated with primary sclerosing cholangitis, a challenging illness with unknown etiology, characterized by chronic bile duct destruction and progression to end-stage liver disease, revealing potential cellular origins of the disease. Our tool, together with this extensive compendium, serves as an invaluable resource for researchers, enhancing our understanding of gene regulation in various tissues and conditions.

## Results

### Deep learning architecture of DeepDETAILS

As the fundamental building blocks of the eukaryotic genome packaging/architecture, the placement of nucleosomes influences many transcriptional regulatory processes. The genome-wide landscape of the nucleosome-depleted regions, also known as open chromatin regions, indicates active regulatory regions that are highly specific to cell types^30–32^. With sc/sn-ATAC-seq becoming increasingly available and affordable for diverse samples, we sought to use chromatin accessibility information for each cell type to reconstruct other regulatory processes from various bulk sequencing assays. To achieve this, we developed DeepDETAILS, an innovative deep learning framework designed to reconstruct cell-type-specific regulatory signals from bulk data (Fig. 1a). DeepDETAILS uses branches of dilated convolutional neural network blocks to make individual predictions for each cell type, which are then combined linearly to reconstruct the bulk signal. While deep learning has previously been used in genomics, most efforts have focused on fully supervised settings that aim to learn regulatory sequence patterns^33^ or predict the functional impact of noncoding variants^34,35^. These models typically rely on direct, experimentally measured signals, such as transcription initiation^36^ or histone modifications^37^, from bulk samples for model training (Fig. 1b). In contrast, DeepDETAILS is uniquely designed for the deconvolution setting, where cell-type-specific signals are not directly available. Instead of learning from ground-truth signals for each cell type, DeepDETAILS operates in a quasi-supervised manner by minimizing the differences between the predicted and actual observed bulk signals (Fig. 1b).

**Fig. 1.**
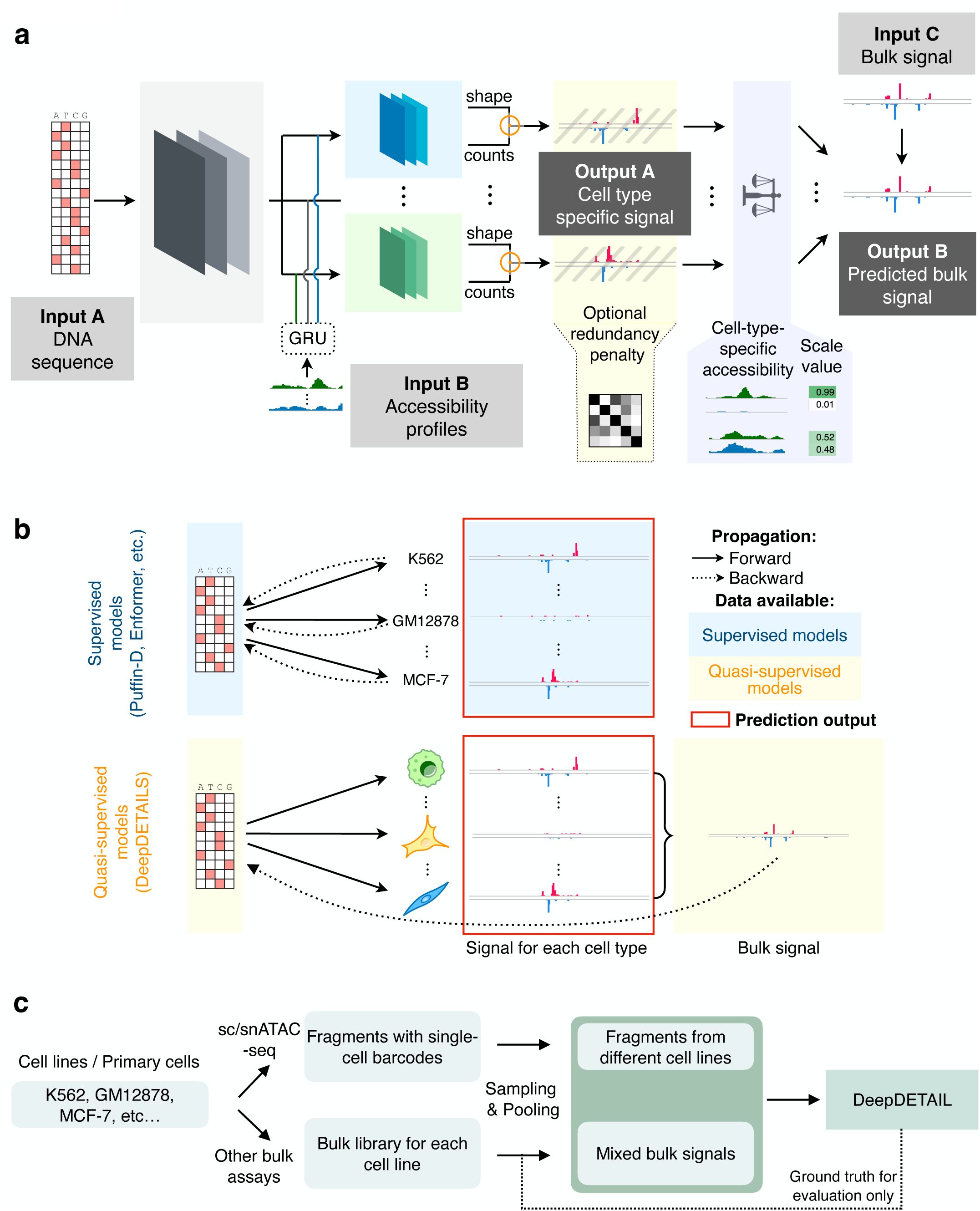
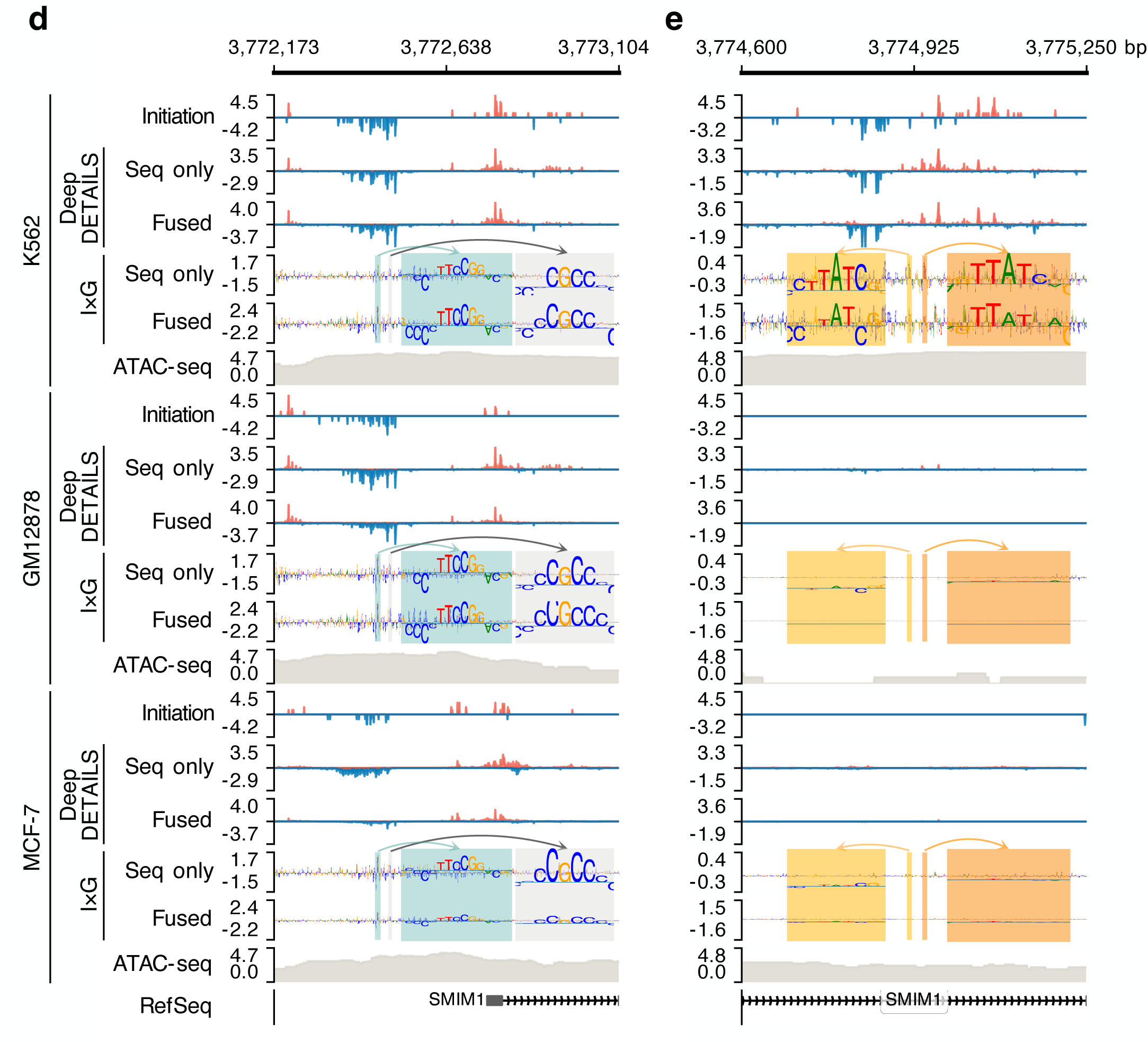
Architecture of DeepDETAILS. **a**, Schematic overview of the DeepDETAILS framework. The model takes three inputs: (i) DNA sequence information, (ii) cell-type-specific accessibility profiles derived from sc/snATAC-seq data, and (iii) ‘bulk’ sequencing signals. DeepDETAILS is optimized to minimize the difference between real and predicted bulk tracks. Note that the “seq only” version of DeepDETAILS does not integrate sequence information with accessibility features extracted by the GRU module (dashed box). **b**, The difference in training approaches between quasi-supervised methods (e.g. DeepDETAILS) and supervised models (i.e. Puffin-D^36^ Enformer^37^, etc.). **c**, Workflow of simulating bulk and reference libraries for evaluating the performance of cross-modality deconvolution. In **d** and **e**, DeepDETAILS deconvolved a simulated bulk PRO-cap library consisting of three cell types: K562, GM12878, and MCF-7. The ground truth signals were labeled as “Initiation”, while deconvolved signals from the “seq only” and “fused” models were shown separately. Sequence motifs contributing to the prediction in each cell type were shown in the “I×G” (input × gradient) tracks. In **d**, motifs known to be important for transcription initiation in general were highlighted. In **e**, motifs associated with cell-type-specific identity were highlighted (e.g., GATA1-like motif for K562, with the consensus sequence GATAA/TTATC). Unless explicitly specified, inverse hyperbolic sine (asinh) transformation was applied to initiation and chromatin accessibility by default to account for their large dynamic range (e.g., between promoters vs enhancers). The asinh transformation is similar to log transformation but supports negative values.

Inspired by the fact that open chromatin is a necessary but not sufficient condition for many regulatory processes (Supplementary Fig. 1), we introduced a constraint condition into DeepDETAILS’ architecture based on the accessibility of the local region, so that if a locus is not accessible in one cell type, tentative predictions made in this cell type will not be incorporated into the final prediction. This forms as a feedback mechanism to the training process and introduces asymmetricity in the architecture.

We implemented two versions of DeepDETAILS (Supplementary Fig. 2). One version used only the DNA sequence and the constraint condition from chromatin accessibility (which we referred to as the “seq only” model), which aims for maximizing the understanding of contribution of DNA sequences. The other version fused the base pair resolution profiles of chromatin accessibility learned by gated recurrent units (GRUs) with the sequence embeddings learned by CNNs (which we referred to as the “fused” model). Fusion of accessibility embeddings allowed for a smaller architecture to extract information from DNA sequences, leading to smaller footprints on GPU memory (Supplementary Fig. 2f). It is also more scalable and flexible, enabling the deconvolution of more cell types given the same configuration of hardware (Supplementary Fig. 2g).

### DeepDETAILS learns TF motifs that govern cell-type agnostic general regulatory processes and cell identity

To validate DeepDETAILS, we conducted a proof-of-concept study using a simulated heterogeneous “tissue” sample composed of three distinct cell types: K562, GM12878, and MCF-7 (Fig. 1c). We generated the bulk library by combining GRO/PRO-cap data, which captures transcription initiation signals from all enhancers and promoters genome-wide, for each cell line, and utilized snATAC-seq of these cell lines as the reference library (Fig. 1c, Methods). When applied to the simulated libraries, both the “seq only” and “fused” versions of DeepDETAILS successfully reconstruct the initiation profiles for all three cell types (Fig. 1d and e).

To interpret the sequence motifs that DeepDETAILS employed in its predictions, we used the input × gradient method^38^. The framework identified motifs associated with transcription initiation in general, including binding sites for Sp/KLF and ETS family of transcription factors (Fig. 1d), which align with findings from previous studies^36,39,40^. Additionally, DeepDETAILS captured cell-type-specific motifs reflective of cell-type identity. For instance, GATA1 is a pioneer transcription factor critical for erythroid differentiation and is notably active in K562 cells^41^. DeepDETAILS learned prominent GATA1-like motifs in K562 cells but not in other cell types (Fig. 1e). While both models produced similar deconvolution results (Fig. 1d and e), the “seq only” model demonstrated better learning of DNA syntax compared to the “fused” model (Fig. 1e). This difference is expected because the “fused” model can partially leverage accessibility embeddings to infer cell type identity and was designed with a smaller architecture for efficiency. Given its hardware-friendly design, we proceeded with the “fused” version for subsequent analyses in this study.

### DeepDETAILS can deconvolve regulatory signals from a wide variety of bulk sequencing **assays**

We assessed DeepDETAILS for its ability to deconvolve regulatory signals from distinct bulk sequencing assays, including transcription initiation as captured by PRO-cap (Fig. 1d, e and Fig 2a), histone modifications as detected by ChIP-seq (H3K27ac, H3K4me1, H3K4me3, Fig. 2b∼d and Supplementary Fig. 3a∼c, respectively), and pause-release dynamics through PRO-seq (Supplementary Fig. 3d). Each assay captures different aspects of regulatory processes at varying resolutions: ChIP-seq provides broader signals, while PRO-cap/seq offers base-pair resolution.

**Fig. 2.**
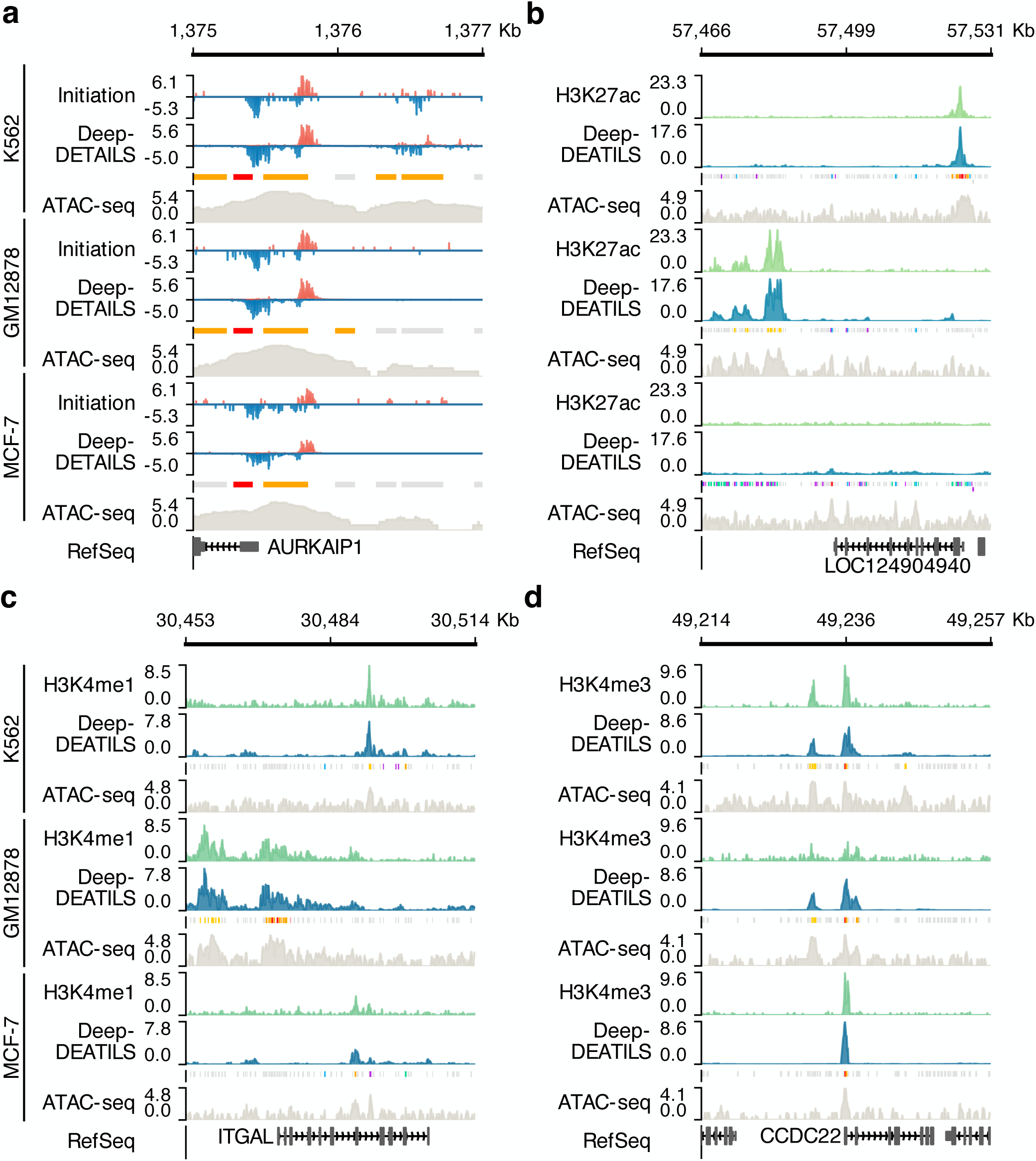
DeepDETAILS works for multiple bulk sequencing assays. **a**, DeepDETAILS deconvolved simulated bulk transcription initiation profiles captured by GRO/PRO-cap from both promoters (labeled in red boxes) and enhancers (labeled in orange boxes). **b**∼**d**, DeepDETAILS also successfully deconvolved bulk histone modification signals, including: **b** for H3K27ac, **c** for H3K4me1, and **d** for H3K4me3. Candidate regulatory elements such as enhancers and promoters were annotated based on ENCODE cCRE definitions^80^.

Despite these differences in assay types and data characteristics, DeepDETAILS consistently performed well across all assays, demonstrating its robustness and versatility. This capability makes it an invaluable tool for analyzing complex biological samples with mixed cell populations.

### Deconvolution by DeepDETAILS matches or exceeds performance of canonical supervised direct-prediction methods

DeepDETAILS operates in a deconvolution setting, where it only has access to bulk signals and not the desired prediction target (cell-type-specific signals). Despite facing a more challenging learning task than previous deep learning studies (Fig. 1b), we evaluated its performance against published fully supervised models for sequence learning^36,37^. The assessment employed another simulation setup where a set of bulk sequencing libraries were created by combining signals from five different cell lines (Methods) that maximized or minimized the pairwise similarity of chromatin accessibility profiles between cell lines when data is available (Supplementary Fig. 4a). This approach aimed to assess how effectively DeepDETAILS could handle samples with varying cell population structures. For example, we simulated two sets of bulk libraries capturing nascent transcription initiation: A673, HCT116, human umbilical vein endothelial cells (HUVEC), K562, and MCF 10A (maximized similarity) and Caco-2, Calu3, GM12878, LNCaP, and MCF-7 (minimized similarity, all simulation formulas are available in Supplementary Table 2). We used DeepDETAILS to reconstruct signal tracks of initiation, pause-release, and histone H3K4me3/H3K27ac modifications for each cell line, and compared its performance with two state-of-the-art supervised models: Puffin-D^36^ and Enformer^37^. Puffin-D was developed for predicting transcription initiation, which we also repurposed to study pause-release by training separate models using the ground truth (Methods). Predicted profiles of histone modifications from Enformer were retrieved from the original study^37^. Since Enformer predicts signals at 128bp resolution, we binned the predictions from both DeepDETAILS and Puffin-D to 128bp to match the resolution. We compared the similarity between the predictions (DeepDETAILS, Puffin-D, and Enformer) and the actual signal tracks for each cell type using Pearson’s *r* as reported in the original studies (Fig. 3a). DeepDETAILS achieved similar performances to state-of-the-art supervised models (Pearson’s *r* for DeepDETAILS: 0.709 (initiation), 0.719 (pause-release), 0.767 (histone modification); other models: 0.726, 0.734, 0.679, respectively) (Fig. 3a). Additionally, since both DeepDETAILS and Puffin-D make predictions at base pair resolution, and the assays for studying transcription initiation and pause-release (PRO-cap/3′ of CoPRO) yield base pair resolution profiles, we tested the performances of the models in recovering the dominant transcription start sites (TSSs) and pausing sites at 1-bp resolution. We demonstrated that DeepDETAILS achieved comparable performance to state-of-the-art models with large receptive fields in predicting dominant TSSs and pausing sites. Specifically, DeepDETAILS obtained Pearson’s *r* values of 0.800 for TSSs and 0.837 for pausing sites, closely matching the 0.817 and 0.836 scores of Puffin-D (Fig. 3b and c; Supplementary Fig. 4b and c). This performance is particularly impressive given that DeepDETAILS was trained on simulated bulk data (mixture of 5 cell lines), whereas Puffin-D and Enformer were trained on ground truth bulk data for each cell line.

**Fig. 3.**
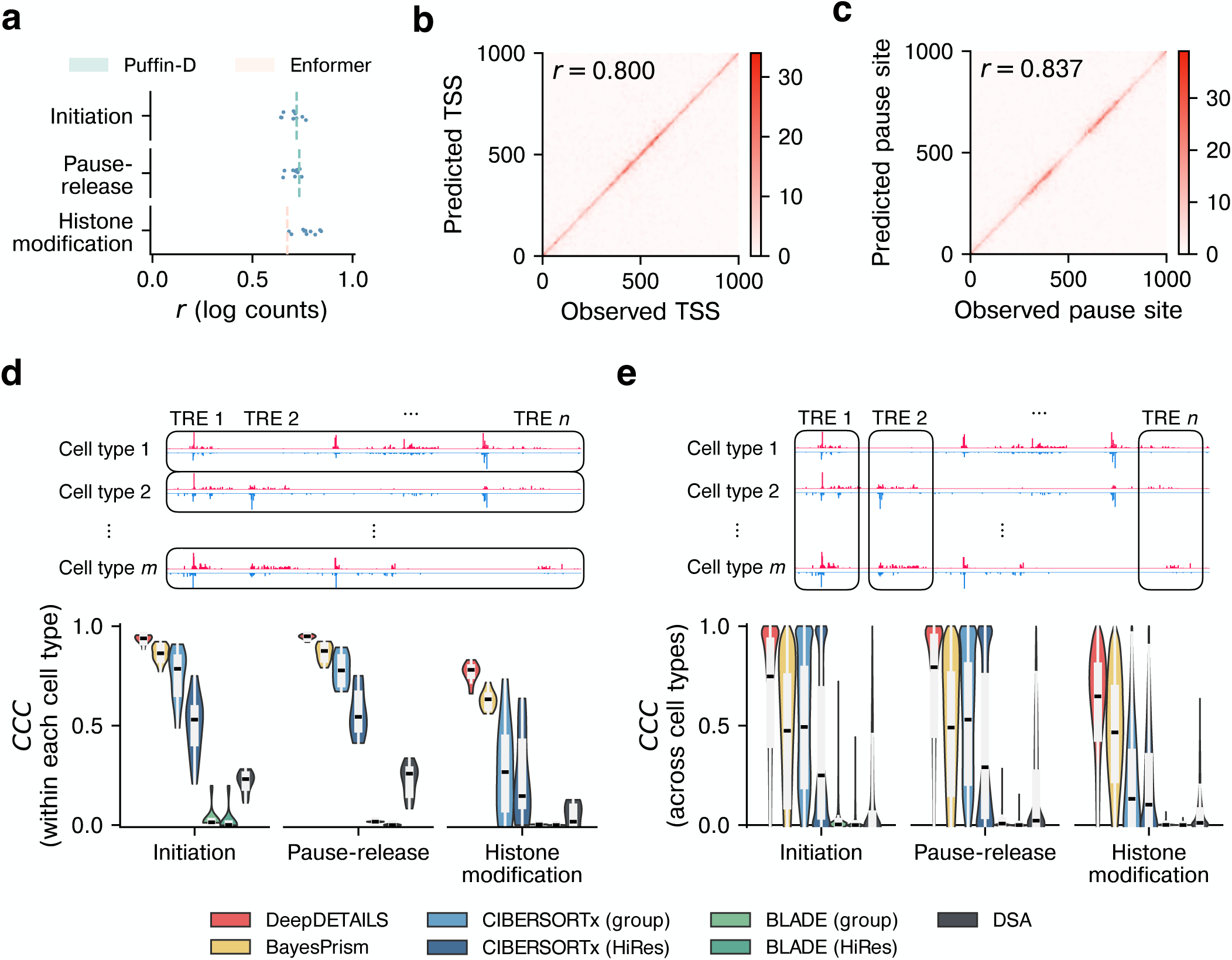
Performance comparison of DeepDETAILS with state-of-the-art methods. **a**, DeepDETAILS was compared against state-of-the-art supervised deep learning models, Puffin-D (for transcription initiation/pause-release) and Enformer (for histone modification). The evaluation used Pearson’s correlation coefficient (*r*) at 128-bp resolution with log-transformed read counts. Dashed line shows the median of *r* that the corresponding supervised model archived. *n* = 10. **b**, Pearson’s correlation coefficient (*r*) was calculated to evaluate consistency between DeepDETAILS’ reconstructed dominant transcription start sites (TSSs) and actual dominant TSSs in the ground truth GRO/PRO-cap libraries. The analysis sampled 5000 peak regions for this assessment. **c**, Similarly, Pearson’s correlation coefficient (*r*) was used to assess consistency between DeepDETAILS’ reconstructed dominant pause sites and actual dominant sites at the 3′ end of the CoPRO libraries. This evaluation also utilized a sample of 5000 peak regions. **d** and **e**, Performance comparison between DeepDETAILS and statistical deconvolution tools (1-kb resolution). **d** for within-each-cell-type evaluation, and **e** for across-cell-type evaluation. Black boxes atop each panel indicate the units used for CCC calculations.

### DeepDETAILS outperforms statistical deconvolution tools in cross-modality settings

Statistical tools are widely used in deconvolving bulk profiles with references of the same modality^22–26^. These tools consider bulk sequencing profiles as linear combinations of signals from each cell type and calculate cell-type-specific signals by solving a system of equations^22–26^. However, the references, which are typically single cell profiles, may not be available for assays that are technically difficult to be performed at single cell resolution (e.g., PRO-cap).

DeepDETAILS addresses this limitation by enabling cross-modality deconvolution. It utilizes references from one modality (sc/snATAC-seq) to deconvolve bulk profiles of another modality (e.g., PRO-cap), thereby overcoming the hurdle of unavailable references for certain assays. Due to the fact that open chromatin regions (i.e. ATAC-seq signals detected) may not have regulatory signal from other processes (e.g. PRO-cap signal detected, Supplementary Fig. 1), canonical statistical methods for same-modality deconvolution may not work for a cross-modality setting, because they operate under the assumption that aggregating reference signals for each cell type can reproduce the bulk data.

To evaluate the performance of DeepDETAILS in cross-modality deconvolution, we compared it with several statistical deconvolution methods – BayesPrism^23^, CIBERSORTx^22^, BLADE^24^, and DSA^42^ – using simulated bulk samples (Fig. 3d and e). Although DeepDETAILS provides predictions at base-pair resolution, statistical tools typically generate count tables for genes or regions. For a fair comparison at the same resolution, we added up predictions from DeepDETAILS for 1 kb regions, and used statistical tools to reconstruct read counts tables in the same 1kb peak regions (Methods). DeepDETAILS demonstrated superior performance compared to statistical methods, as expected (Fig. 3d and e). It achieved the highest Lin’s concordance coefficients (CCC) when comparing values for peaks within each cell type: 0.938 for initiation, 0.949 for pause-release, and 0.780 for histone modifications. The best-performing statistical methods yielded CCCs of 0.864, 0.876, and 0.632, respectively (Fig. 3d). Similarly, DeepDETAILS excelled when comparing peak values across different cell types with CCCs of 0.748, 0.793, and 0.647 for initiation, pause-release, and histone modifications, outperforming the statistical methods’ results of 0.493, 0.530, and 0.466 (Fig. 3e and Supplementary Fig. 4d). In addition to evaluating how closely predicted values align with observed ones through a correlation metric (CCC), we also assessed the precision of our predictions by calculating an error metric (root mean squared error, RMSE). DeepDETAILS reconstructed read counts for each cell type with highest accuracy compared to other methods, as evidenced by mean overall RMSEs of 0.947 (Supplementary Fig. 4e). Additionally, statistical deconvolution methods usually require multiple bulk samples^22–24,42^ or reference samples^25^, DeepDETAILS only need one bulk and reference sample, which enables its broader applications in research projects of different scales and purposes.

### DeepDETAILS works robustly in complex settings

Many factors can significantly impact the accuracy and reliability of deconvolution outcomes. Therefore, we aimed to evaluate the performance of DeepDETAILS under more demanding conditions. First, we examined the robustness of DeepDETAILS when applied to libraries with varying sequencing depths. We simulated bulk and reference libraries across three distinct sequencing depth categories: excellent, good, and acceptable (Methods). DeepDETAILS maintained consistent performance levels, with minimal reductions observed as the sequencing depths of the libraries decreased. Specifically, using CCC as a metric, within-cell-type comparisons demonstrated limited performance decline when transitioning from excellent sequencing quality to good quality (range of decreasing: 0.004 to 0.025, Fig. 4a), and from good to acceptable quality (range of decreasing: 0.034 to 0.097, Fig. 4a). Similar trends were observed for across-cell-type comparisons (Supplementary Fig. 5a). These results indicate that DeepDETAILS exhibits robust performance across different sequencing depths, making it a reliable tool for deconvolution analysis even under suboptimal conditions.

**Fig. 4.**
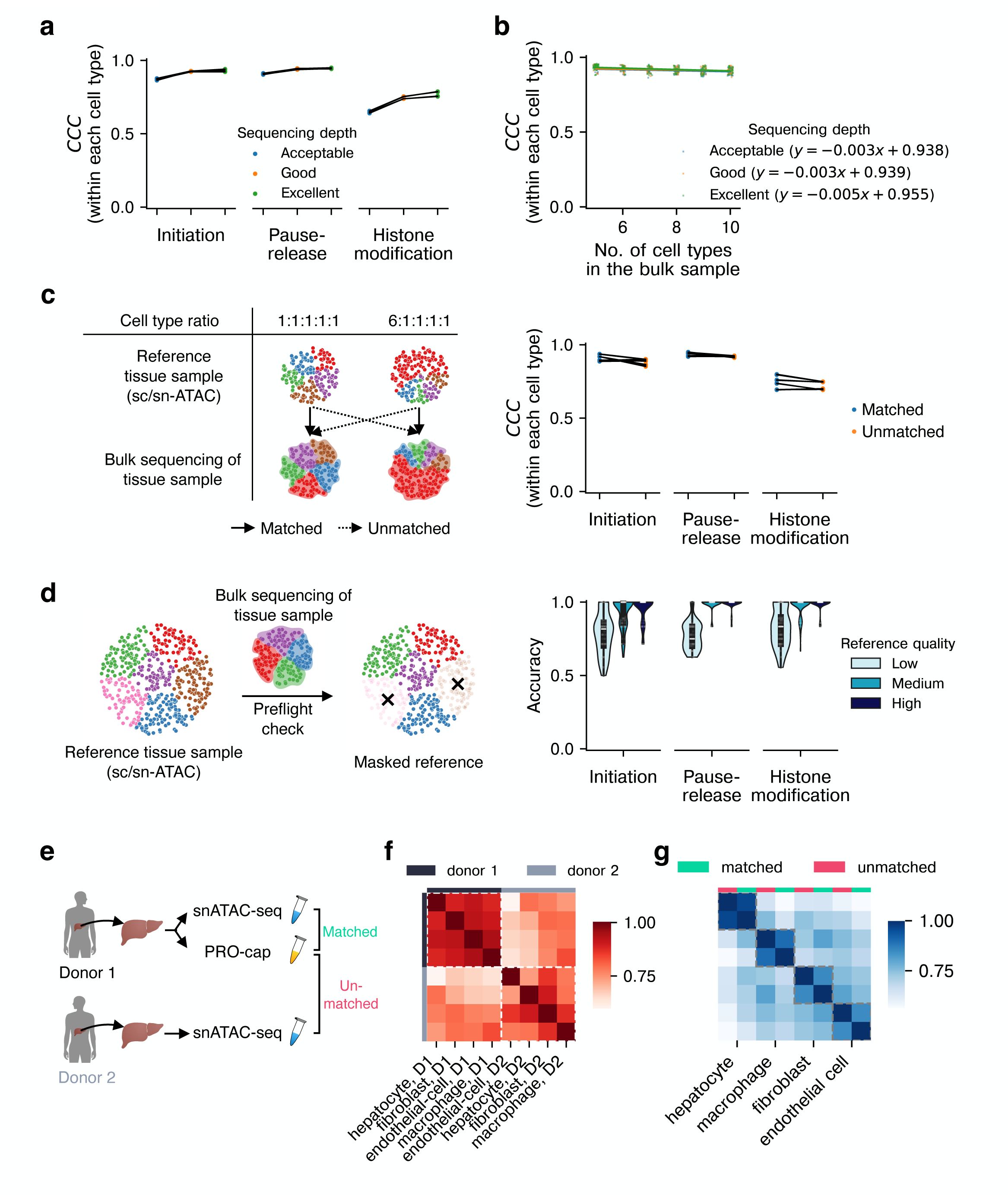
Performance evaluation of DeepDETAILS under challenging cases. **a**, DeepDETAILS’ performance by varying the sequencing depths of bulk and reference libraries. For each sequencing technique, two sets of bulk-reference pairs were simulated, and deconvolution was repeated three times (*n* = 6 per technique). **b**, Evaluation of DeepDETAILS’ performance on deconvolving bulk transcription initiation sequencing libraries with different numbers of cell types in the bulk and reference (*n* = 3 × *k* for a simulated setting with *k* cell types). **c**, Schematic (left) and results (right) of using DeepDETAILS to deconvolve bulk samples with unmatched reference libraries (ratios of cell types do not match in the bulk and reference, *n* = 6). For **a** ∼ **c**, evaluation was done by calculating the CCC within each cell type as done in Fig. 3d. **d**, Schematic of the preflight check step (left) and the accuracy of preflight in masking out cell types in the reference but not in the bulk (right, *n* = 91 for each condition). **e**, Schematic of evaluating DeepDETAILS with real datasets. **f**, Genome-wide correlation analysis (Pearson’s *r*) suggested the pairwise similarity relationships between different cell types were disrupted in the two reference samples. **g**, The deconvolved signals from DeepDETAILS maintained the pairwise similarity (Pearson’s *r*) relationships regardless of the choice of reference libraries.

Cell type composition in human tissues can be highly complex; therefore, a key attribute of deconvolution methods is their capacity to effectively accommodate this variation in heterogeneity. We simulated bulk libraries containing 5 to 10 cell types and assessed DeepDETAILS’ performance across three sequencing depth conditions. We found that the number of cell types in the bulk library has only a minimal impact on both within-and across-cell-type CCCs. Specifically, the inclusion of one additional cell type results in a decrease of 0.003 in within-cell-type CCC (Fig. 4b) and 0.010 in across-cell-type CCC (Supplementary Fig. 5b).

Previous simulations assumed that the bulk and reference samples originated from the same source and shared identical cell type ratios. Expanding DeepDETAILS’ ability to work with unmatched samples—i.e., those from different sources or with varying cell type proportions— would significantly enhance its practicality. To evaluate DeepDETAILS’ capability, we simulated two distinct groups of samples: one group featuring an even distribution of cell types and another dominated by a single cell type (comprising 60% of the total). “matched deconvolution” was defined as using reference and bulk samples from the same group, while “unmatched deconvolution” involved pairing samples from different groups (Fig. 4c). The simulated unmatched settings were designed to represent extreme scenarios with large fluctuations in cell type ratios. However, it is important to note that typical fluctuations across different human tissue samples are usually small (Supplementary Fig. 5c) as indicated in the ENCODE snATAC-seq experiments set^3^. By demonstrating robust performance under these extreme conditions, we validate DeepDETAILS’ ability to handle the more common small fluctuations as well. Our results showed consistent deconvolution outcomes in donor matched and unmatched settings, with differences in within-and across-cell-type CCCs ranging from 0.014 to 0.026 and 0.031 to 0.082, respectively (Fig. 4c and Supplementary Fig. 5d).

Consortium-level efforts^3,43^ have been increasing the availability of human tissue sc/snATAC-seq libraries, so users may not need to generate their own reference libraries to deconvolve their bulk samples with DeepDETAILS. Reference sc/snATAC-seq libraries provided by consortium are usually generated by pooling multiple specimens to ensure a comprehensive representation of different cell types, so there may be more cell types in the reference libraries than the bulk sample. To address this, DeepDETAILS employs a “preflight check” step that estimates the proportions of different cell types within the bulk sample. It further identifies and masks out cell types present in the reference but absent in the bulk, preventing their influence on subsequent analyses (Methods). We evaluated the accuracy of the preflight step by blending in 1-5 more additional cell types into the references used by DeepDETAILS. The results demonstrated high accuracy (median accuracy of 1.0) in detecting extraneous cell types when the reference libraries had adequate sequencing depth (Fig. 4d).

Finally, to validate the robustness of DeepDETAILS for real-world applications, we applied this method to reconstruct cell-type-specific initiation maps using a bulk liver PRO-cap library (Fig. 4e). We utilized a matched snATAC-seq library from the same donor’s right lobe of the liver^44^. To further assess its performance in unmatched conditions, we deconvolved the same bulk library using another snATAC-seq dataset derived from a different donor’s liver (Fig. 4e). We observed significant batch effects between the two snATAC-seq libraries, with accessibility tracks clustering by their sources rather than cell type identities (Fig. 4f and Supplementary Fig. 5e). Additionally, there were noticeable disruptions in pairwise similarities between cell types across donors. For instance, hepatocytes and endothelial cells showed high similarity (Pearson’s *r* = 0.911) in one donor, whereas fibroblasts displayed strong similarity to hepatocytes (Pearson’s *r* = 0.896) in another (Fig. 4f). Despite these challenges, DeepDETAILS demonstrated remarkable resilience by reconstructing consistent tracks across both matched and unmatched settings. The method achieved strong correlations for all cell types (Fig. 4g): hepatocytes (Pearson’s *r* = 0.959), macrophages (*r* = 0.898), fibroblasts (*r* = 0.849), and endothelial cells (*r* = 0.846). This robust performance underscores DeepDETAILS’ capability to handle batch effects and donor variability effectively, ensuring reliable results in real-world scenarios.

### A cell-type-specific compendium of regulatory maps in human tissues

Projects such as the NIH Roadmap Epigenomics Program^45^ and ENCODE^3^ have significantly advanced our understanding of gene regulation across various human tissues and developmental stages. However, these initiatives primarily utilized bulk sequencing techniques, which report signals from mixed cell populations rather than focusing on individual cell types. To address this limitation, we developed a comprehensive resource comprising regulatory maps specific to different cell types across the human body. This resource enhances our ability to understand transcription regulation, annotate noncoding regions of the genome, and prioritize noncoding genetic variants associated with diseases. Using DeepDETAILS, we deconvolved bulk ChIP-seq datasets from the ENCODE project, focusing on key histone modifications including H3K4me1, H3K4me3, and H3K27ac, in tissues for which snATAC-seq references were available.

Additionally, we generated genome-wide maps for transcription initiation at promoters and enhancers across 10 human tissues and applied DeepDETAILS to deconvolve their cell-type-specific components. (Fig. 5a and Supplementary Fig. 6a). The resulting compendium includes massive regulatory profiles across 20 organs, 39 tissues, and 86 distinct cell types. All datasets are publicly available through our web portal (https://details.yulab.org), where users can explore the deconvolved results by assay type and biosample category. The portal also includes an interactive genome browser for rapid visualization of signal distributions across cell types, along with downloadable files for custom downstream analysis.

**Fig. 5.**
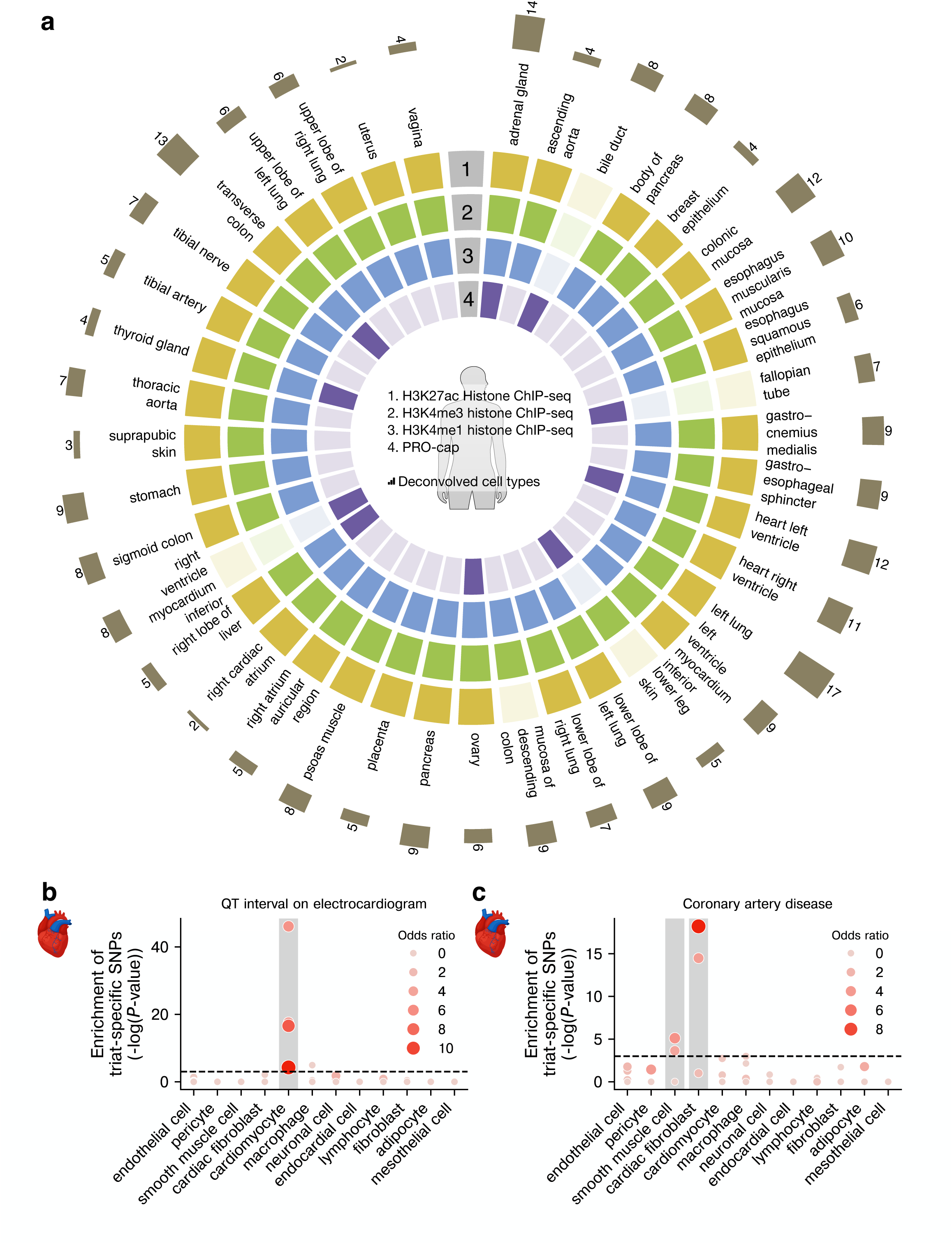
DeepDETAILS compendium. **a**, Summary of DeepDETAILS compendium. Solid color indicates tissues for which there is at least one corresponding bulk sequencing library available in our compendium. Number of dissected cell types in each tissue was labeled in the corresponding bars. **b**, Variants associated with the trait QT interval are enriched in genomic loci that are specifically active in cardiomyocytes. **c**, Variants associated with the disease coronary artery disease are enriched in genomic loci that are specifically active in cardiac fibroblasts and smooth muscle cells.

To demonstrate the quality of our compendium, we identified regulatory elements exhibiting cell-type-specific activity within individual organs (Methods). These elements were enriched for genetic variants associated with specific biological functions or disease, aligning with the known roles of the cell types. For instance, in the heart, cardiomyocytes directly influence the electrical properties and ion channel functions that regulate the duration of the QT interval during repolarization^46^. Our analysis revealed an enrichment of variants associated with these processes specifically in cardiomyocytes, but not in other cardiac cells (Fig. 5b). Cardiac fibroblasts and heart smooth muscle cells play key roles in vascular remodeling, plaque formation, and fibrosis, all of which contribute to the development and progression of coronary artery disease^47,48^. Trends of enrichments were observed specifically in these cell types but not in cardiomyocytes (Fig. 5c). Similar cell-type-specific enrichments were detected for other tissues, including hepatocytes in liver related to complement factor B levels and T-cells in lung concerning COVID-19 severity (Supplementary Fig. 6b and c). To further assess the accuracy of the deconvolved tracks, we also validated them against 98 known cell-type-specific markers^49,50^ for 26 cell types. We observed on average a 17-fold enrichment of signals in the promoter regions of cell-type-specific markers in the corresponding deconvolved tracks (Supplementary Fig. 6d). All these suggest that DeepDETAILS accurately reconstructed signals specific to each cell type.

### DeepDETAILS compendium facilitates the understanding of disease etiology

High-throughput sequencing has revolutionized our ability to study the etiology of various diseases, providing unprecedented insights into genetic and molecular mechanisms^51^. However, analyzing bulk libraries presents unique challenges due to the complexity of signals arising from diverse cell types within tissues. This mix of signals can obscure meaningful patterns, leading to potential misattribution or overlooking critical dysregulations in specific cell populations. DeepDETAILS compendium addresses these challenges by enabling researchers to dissect regulatory signals at cell-type level within bulk samples, thereby enhancing our understanding of disease etiology.

Primary sclerosing cholangitis is a liver disease characterized by chronic inflammation and bile duct scarring, ultimately leading to liver damage and failure (Fig. 6a). However, the underlying cause of the inflammation remains poorly understood. Genome-wide association studies (GWAS) have identified over 300 variants associated with PSC^52–54^; however, pinpointing causal variants remains challenging, in part due to the complex structures of linkage disequilibrium^55^.

**Fig. 6.**
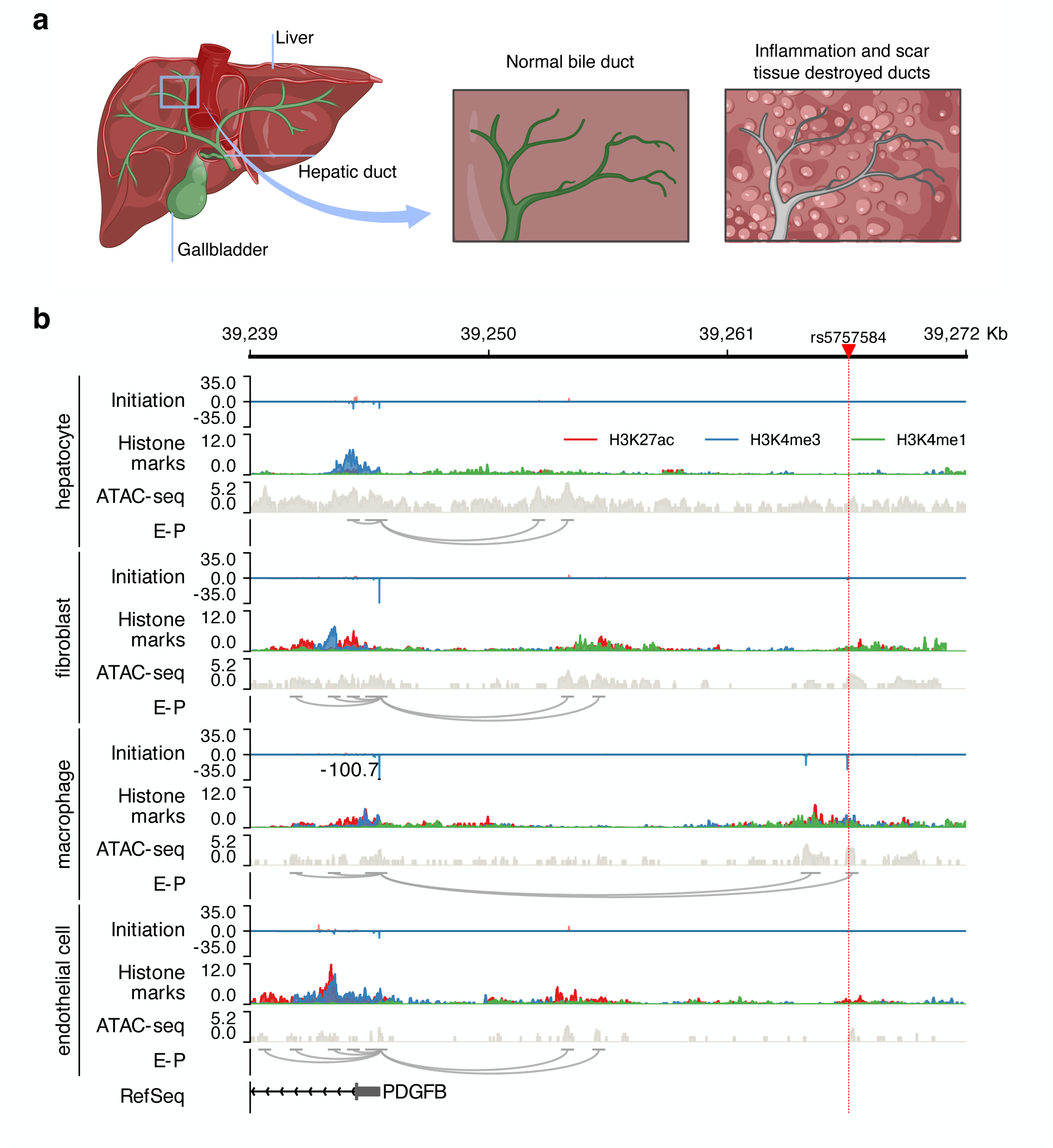
DeepDETAILS compendium brings new insight into disease etiology. **a**, Schematic of primary sclerosing cholangitis. Figure created with BioRender. **b**, Genome browser view of deconvolved signals at the PDGFB locus in the right lobe of human liver. The variant affecting the macrophage-specific enhancer is highlighted at the top of the panel, with the disrupted E-P interaction marked in red. Tracks for transcription initiation and histone modifications are displayed in their original scale.

To better understand the molecular mechanisms underlying PSC, we utilized the reconstructed cell-type-specific regulatory maps in DeepDETAILS compendium for transcription initiation, H3K4me1, H3K4me3, and H3K27ac modifications in the human liver’s right lobe. We then generated cell-type-specific enhancer-promoter interaction (E-P) networks using the activity-by-contact model^56^ with deconvolved histone modification signals as inputs. The predicted E-P interaction maps were refined by removing enhancer candidates lacking deconvolved eRNA transcription. Through this approach, we revisited GWAS-identified PSC-associated variants and identified rs5757584, located within an enhancer that is specifically active in liver-resident macrophages (Kupffer cell) (Fig. 6b). The inferred E-P interaction map suggested that this enhancer regulates a nearby gene, platelet-derived growth factor subunit B (*PDGFB*). To validate this finding, we compared the transcriptional initiation activity of *PDGFB* across cell types to its expression profile in published liver scRNA-seq data^50^, observing high concordance between our results and the scRNA-seq dataset (Supplementary Fig. 7a). Furthermore, this E-P interaction was also supported by an independent eQTL study^57^, which demonstrated that the C-to-T mutation negatively impacted the expression of *PDGFB* in lymphocytes (Supplementary Fig. 7b). Previous studies have shown that the secretion of PDGF from macrophages can help suppress the inflammatory response^58,59^. Therefore, one potential etiology of PSC could be that individuals carrying alternative alleles at rs5757584 exhibit impaired PDGFB-mediated immune regulation in Kupffer cells, resulting in a prolonged inflammatory response and increased susceptibility to liver damage. Notably, hepatocytes account for approximately 70% of liver’s regulatory signals (Supplementary Fig. 7c), suggesting that cell-type-specific effects in non-hepatocytes populations like Kupffer cells could be easily overlooked in bulk-level analyses (Supplementary Fig. 7d). Without the cell-type-specific dissection of regulatory processes provided by DeepDETAILS, the functional role of this variant, both in terms of gene regulation and disease etiology, might have remained undetected (Supplementary Fig. 7d).

## Discussion

Various high-throughput sequencing assays have significantly advanced our understanding of transcriptional regulation. However, it is difficult to use bulk sequencing assays to identify distinct regulatory patterns in tissue samples composed of diverse cell types. While advances in single-cell sequencing (e.g., scRNA-seq, snATAC-seq) allow high-resolution study of regulatory layers in complex biosamples, numerous other assays are still not applicable at the single-cell level. Current machine learning techniques that leverage single-cell sequencing data focused on smoothing technical noises in sc/snATAC-seq sequencing libraries and better representing the clustering structures of the cell population in the sample^60–64^. When data from another single-cell modality is available, such as scRNA-seq, cross-modality data imputation can be performed to further refine the representations^65–68^.

Our DeepDETAILS utilizes the sc/snATAC-seq for a completely different goal: DeepDETAILS reconstructs regulatory processes in different cell types at base pair resolution in a bulk sequencing sample when provided with a sc/snATAC-seq reference. We showed the broad applicability and robust performance of our method in such a cross-modality deconvolution setting. One of the most important applications of DeepDETAILS is its ability to link genetic variants to specific cell types and regulatory pathways, offering new insights into disease mechanisms. For example, the identification of rs5757584 as a variant in a macrophage-specific enhancer affecting PDGFB expression highlights how this tool can pinpoint cell-type-specific functional elements that potentially contribute to disease pathogenesis. This finding underscores the importance of studying regulatory variation at cell-type-specific level and demonstrates DeepDETAILS’ potential to bridge gaps in our understanding of genotype-phenotype relationships.

We further applied DeepDETAILS to deconvolve bulk sequencing libraries across 39 different human tissues and generated cell-type-specific regulatory maps regarding transcription initiation and histone modifications (H3K4me1, H3K4me3, and H3K27ac). We anticipate the comprehensive and high-resolution cell-type-specific regulatory maps in our portal can help identify molecular mechanisms underlying various pathologies, offering new avenues for therapeutic intervention.

While DeepDETAILS represents a leap forward, it is not without limitations. The tool’s effectiveness depends on the availability and quality of sc/snATAC-seq reference dataset. Additionally, the current implementation of DeepDETAILS is not optimized for repressive regulatory processes such as the tri-methylation of lysine 27 on histone H3 protein, and modifications on the architecture may be needed. In addition, statistical deconvolution methods usually benefit from a larger number of input bulk samples, so it is possible that these tools may perform better when more data is available.

Multi-modal foundation models demonstrate potential in generalizing to unobserved cell types and could potentially be repurposed for tasks like deconvolution^69,70^. However, these models require extensive sequencing data and significant computational resources; for instance, EpiBERT needed around 740 ATAC-seq datasets. Their effectiveness also depends on the similarity between regulatory motifs in trained and novel cell types. In contrast, DeepDETAILS’ broad applicability across various cell types, regardless of their similarity, enhances its utility in advanced genomic analyses without the need for vast datasets or powerful computing infrastructure.

In conclusion, DeepDETAILS represents a new advance in the field of genomics, offering researchers a novel framework for dissecting cell-type-specific regulatory landscapes from bulk sequencing data. By enabling fine-grained analysis of transcription regulation and linking genetic variation to disease mechanisms, this tool has the potential to accelerate our understanding of complex traits and inform new approaches to disease treatment.

## Acknowledgements

We thank Nathaniel D. Tippens, Jin Liang, Le Li, Dapeng Xiong, Adam He, Yutong Zhu, and Hongya Zhu for their helpful discussions. We also thank the ENCODE single cell working group and the DCC, especially Austin Wang, Benjamin Doughty, Benjamin Parks, Elisabeth Rebboah, Eran Kotler, Julia Marie Schaepe, Salil Deshpande, Winston Becker, Yu Fu, Wenpin Hou, and Anshul Kundaje, for all their efforts on annotating the cell types. This work was supported by grants from the National Institutes of Health (R01AG077899 and R01HG012970 to J.T.L. and H.Y.). This work used Delta at National Center for Supercomputing Applications and Jetstream2 at Indiana University and Cornell University through allocations BIO240120 and BIO220060 from the Advanced Cyberinfrastructure Coordination Ecosystem: Services & Support (ACCESS) program, which is supported by National Science Foundation grants #2138259, #2138286, #2138307, #2137603, and #2138296.

## Author Contributions Statement

Conceptualization: H.Y.; Methodology: L.Y., J.T.L., and H.Y.; Software: L.Y.; Formal analysis: L.Y. and H.Y.; Investigation: L.Y. and A.O.; Data curation: L.Y., T.X., M.W., and A.K.Y.L.; Resources: S.R.S., J.T.L., and H.Y.; Writing – Original Draft: L.Y.; Writing – Review & Editing: A.O., S.R.S., J.Z., J.T.L., and H.Y.; Visualization: L.Y., X.P., T.X., V.D.F., and H.Y.; Supervision: J.T.L., and H.Y.

## Competing Interests Statement

S.R.S. is an equity holder at OncoVisio, Inc., an equity holder and member of the scientific advisory board at NeuScience, Inc., and a consultant for Third Bridge Group Limited, none of which are related to this work. The remaining authors declare no competing interests.

## Methods

### Processing sequencing libraries

Single-nucleus ATAC-seq libraries for different cell lines and primary cells were processed with 10x Genomics Cell Ranger ATAC 2.1.0 and only fragments from called cells were kept for downstream usage. Run-on sequencing libraries (GRO-cap, PRO-cap, CoPRO, and PRO-seq) data sets were processed with pipelines introduced in our previous publication^11^ and the protocol is available at our GitHub repository. Histone ChIP-seq libraries were processed with the ENCODE ChIP-seq pipeline version 2. All sequencing data and analytical pipelines were managed by BioQueue^71^.

### Simulating bulk tissue datasets for different targeting assays

For run-on libraries (GRO-cap, PRO-cap, CoPRO, and PRO-seq) where only the ends of sequencing reads were considered in downstream analysis, we sampled signals from the corresponding bigWig files with a binomial sampler, so that the sampled signal at locus *i* for cell line *k* follows *s_ik_* ∼ ℬ(*N_ik_*, *p_i_*), where *N_ik_* is the observed signal value from the original library, and *p_i_* is the sampling ratio for cell line *k*. For histone modification ChIP-seq where the entire reads were considered, we downsampled the aligned reads in the tagAlign files following a uniform distribution *u(p_i_*). In both cases, the sampled signals for each cell line were merged together and served as the bulk tissue signal tracks. Peaks were called from the simulated bulk libraries using PINTS^11^, dREG^72^, and MACS2^73^ for GRO/PRO-cap, 3′ of CoPRO/PRO-seq, and ChIP-seq, respectively. We surveyed published run-on and histone modification libraries, and used the 25^th^, 50^th^, and 75^th^ percentiles of their sequencing depth as the criteria of acceptable, good, and excellent simulated bulk library (Supplementary Table 3 for the cutoff for each assay).

### Simulating pseudo-bulk sc/snATAC-seq datasets

We used a two-stage hierarchical sampling strategy to simulate sc/snATAC-seq libraries for tissues. In the first stage, we uniformly sampled *m*_k_ single cells based on their barcodes from the corresponding sc/snATAC-seq library for that cell line / primary cell (*k* in this case); in the second stage, for each sampled cell *c_i_*, we randomly selected *n_i_* reads from the fragments pool of that cell, where *n_i_* ∼ *N*(μ, σ). We used the (μ, σ) pairs (10000, 833), (5000, 333), and (1000, 167) to simulate excellent, good, and acceptable quality sc/snATAC-seq libraries as suggested in a previous study^74^, respectively. The final pseudo-bulk sc/snATAC-seq libraries were obtained by pooling all sampled fragments together. Sampling procedures in both stages were done without replacement.

### DeepDETAILS

The expanded architectural specifications of DeepDETAILS, including the number of layers, channels, filter sizes, and activation functions, are presented in Supplementary Fig. 2. At its core, both versions of DeepDETAILS feature bodies composed of dilated CNN layers designed to learn embedding from the 4096 bp input DNA sequence. These embeddings are passed to *K* pairs of heads for predicting the shape of the signals and the absolute counts in each cell type for the center 1000 bp window (*K* is the number of cell types in the bulk sample), similar to a previous study^75^. The predictions for each cell type are adjusted using a scale function that zeros out predictions from an inaccessible region in specific cell types. The scaled predictions from individual cell types are aggregated to form the predicted bulk signal. Both peak regions identified from the bulk libraries as well as background regions with similar GC content distributions will be used for training.

Scale function for cell type *k* at region *r*:

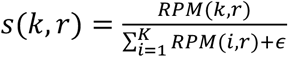

where RPM(*k, r*) gives the depth-normalized ATAC-seq signals at the center 1-kb region of *r* in the corresponding cell type *k*, and ϵ = 10^-16^ to avoid dividing by zeros.

During training, DeepDETAILS minimizes the root mean squared logarithmic error (RMSLE) between the predicted and observed signals. To prevent the model from collapsing into trivial solutions where multiple branches predict similar outputs, DeepDETAILS implements a redundancy penalty^76^. This penalty discourages excessive correlation between predictions from different cell types and improves the overall robustness of the model. Mathematically, the loss function can be expressed as:

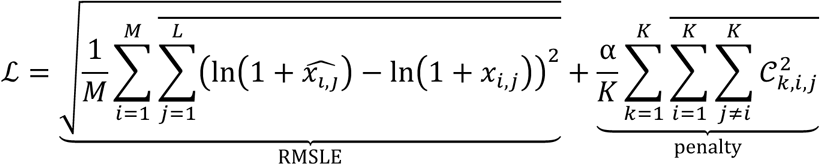

where *M* is the number of regions in the batch, *L* is the length of the prediction window, *K* is the number of cell types, α is a positive constant to balance the RMSLE and the penalty, and *c* is the correlation matrix of predictions.

### Comparing DeepDETAILS with supervised learning methods

We trained two sets of Puffin-D^36^ models, and each model was fed by downsampled signals from five sources (cell types) as the authors did in their original manuscript. In one setting, Puffin-D models were trained to predict signals for transcription initiation and pause-release in the other.

The combinations of cell lines were identical to what we used in our simulated bulk samples for testing the performance of DeepDETAILS and the formulas are available in Supplementary Table 2. Predictions of Enformer were retrieved from the original publication^37^ and only predictions for biosamples that were also tested in this study were kept for comparison.

Comparisons were evaluated using chromosomal cross validation.

### Benchmark performances on cross-modality deconvolution for statistical methods

Statistical deconvolution methods were used to purify total read counts among regions of interest in 1-kb resolution for each mixed cell type. For CIBERSORTx^22^, we applied S-mode batch correction and lowered the filtering threshold for average gene expression to 0, as suggested in the original publication. For BayesPrism^23^, BLADE^24^, and DSA^42^, we used the default parameters. Invalid predictions (not a number) were replaced by small random values drawn from a distribution that ln(*X*) ∼ *N*(0,1) and scaled by 0.1, when calculating correlation coefficients between the predicted values and the groundtruth. Additional bulk samples were simulated since statistical methods require multiple bulk samples to run.

### Preflight check

When preparing the data files for running DeepDETAILS’ deconvolution workflow, a preflight procedure will be executed by default. This step trims the reference dataset by only keeping cell types consistently present in both the reference and bulk datasets. We implemented this functionality in a three-steps manner: first, we build a signature matrix as previously mentioned^22^, then we use *v*-SVR to estimate the fractions of different cell types in the bulk library as suggested by a previous study^22^, lastly cell types with very low presence are removed (<3.5%) unless the preflight check is manually bypassed with option (--skip-preflight). While this method may yield lower accuracy in estimating the absolute fraction (median of *R*^3^: 0.567∼0.827) compared to intra-modality simulations^77^, it remains effective for identifying cell types absent from the bulk sequencing library.

### Generation of PRO-cap libraries for human tissues

Human tissue samples were obtained as part of the ENCODE consortium’s coordination effort. PRO-cap library preparations for two biological replicates, each containing 10 million cells, were performed separately as previously described^5,78,79^. In brief, cells were permeabilized, followed by run-on reactions. RNA was then isolated, and two rounds of adaptor ligation and reverse transcription were carried out using custom adaptors. 5′ cap selection was performed between adaptor ligations through a series of enzymatic steps. RNA was washed followed by phenol:chloroform extractions, and ethanol precipitations between reactions, all under RNase-free conditions. After PCR amplification and library clean-up, libraries were sequenced on an Illumina NovaSeq lane.

### Deconvolution of human tissue samples

For the deconvolution of bulk human tissue samples, we utilized DeepDETAILS with its default settings. For reference dataset selection, we prioritized single-nucleus ATAC-seq (snATAC-seq) data derived from the same tissue and donor. In cases where such data were unavailable, a composite reference was constructed using all available snATAC-seq datasets for that tissue. The specific reference dataset used for each deconvolution experiment is documented on our web portal. Each deconvolution workflow was performed in duplicate, with average signal values being reported to ensure robustness of the results. For histone ChIP-seq experiments, we merged the deconvolved signals on the two strands and reported binned signals (10 bp) to the portal.

### Identification of cell-type-specific regulatory elements and variant enrichment analysis

We identified candidate regulatory elements exhibiting cell-type-specific activity in different tissues/organs using *t*-tests, by comparing the depth-normalized signal of a specific assay in a given cell type to that of all other cell types. We controlled for false discovery rate using Benjamini-Hochberg’s approach and considered regions with an FDR < 0.05 and log fold change > 2 as significant. To assess enrichment of variants, we applied Fisher’s exact test for each trait. Within each organ, we identified GWAS variants located in bulk peak regions and categorized them into two groups: those residing in cell-type-specific regions and those in non-specific regions. We then further classified these variants based on their association with a specific trait and constructed 2 × 2 contingency tables to facilitate odds ratio calculations.

Finally, for each cell type, we calculated the odds ratio of trait-associated variants occurring in cell-type-specific regions compared to non-specific regions and reported the results. Trait-cell-type pairs with adjusted *p*-values (Benjamini-Hochberg) smaller than 0.05 were considered significant, indicating a statistically significant enrichment of trait-associated variants in cell-type-specific regions.

### Prediction of enhancer-promoter interaction

Activity-by-contact model was used for predicting E-P interactions^56^. For the cell-type specific E-P maps, we used pseudo-bulk ATAC-seq track for the cell and deconvolved H3K27ac ChIP-seq signal as the measurement of enhancer activity. For the bulk E-P map, we used the entire pseudo-bulk ATAC-seq and bulk H3K27ac ChIP-seq as the enhancer activity measurement. In both cases, we used Hi-C tracks averaged from multiple human biosamples as the measurement of contact frequency. Predicted interactions were further pruned by removing pairs where the candidate enhancer anchors do not have eRNA transcription (less than 20 counts per million reads).

### Code availability

The source code of DeepDETAILS is publicly available at https://github.com/haiyuan-yu-lab/DeepDETAILS, scripts used to generate results that are reported in this study can be retrieved from https://github.com/haiyuan-yu-lab/DeepDETAILS-analysis.

### Data availability

All deconvolved tracks are freely available on our portal. Sequencing data retrieved from public databases (NCBI GEO and ENCODE portal) are listed in Supplementary Tables 1. Cell type annotations for sc/snATAC-seq experiments were obtained from the ENCODE portal and the corresponding accession for each deconvolution run is available on our online portal. The data used for the eQTL analysis described in this manuscript were obtained from the GTEx Portal on 12/13/2024.

## Notes

### Competing Interest Statement

The authors have declared no competing interest.

