## Supplementary figures 1~7 for "High-resolution reconstruction of cell-type specific transcriptional regulatory processes from bulk sequencing samples"

#### Supplementary Fig. 1

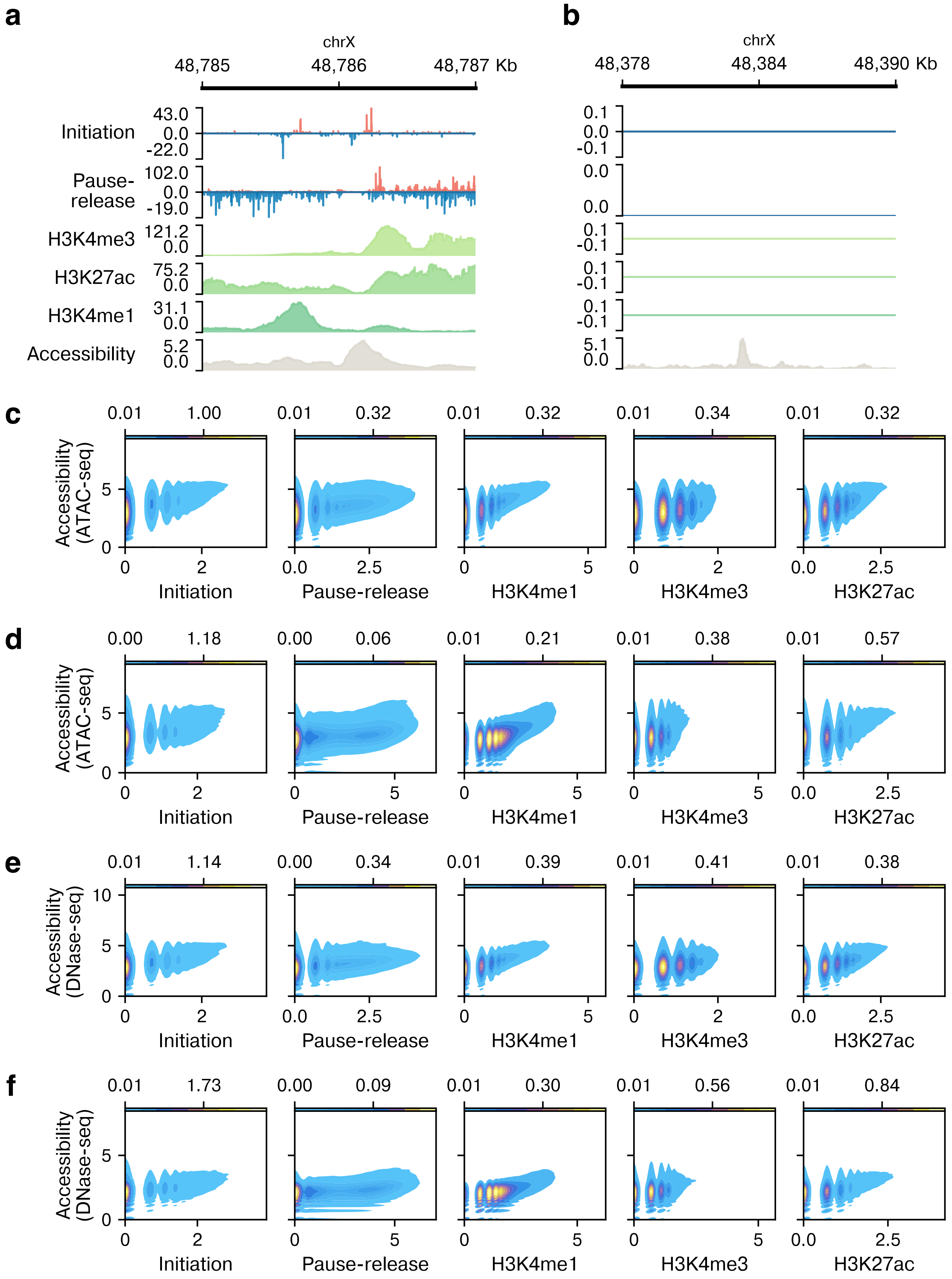

### Supplementary Fig. 2

**a**

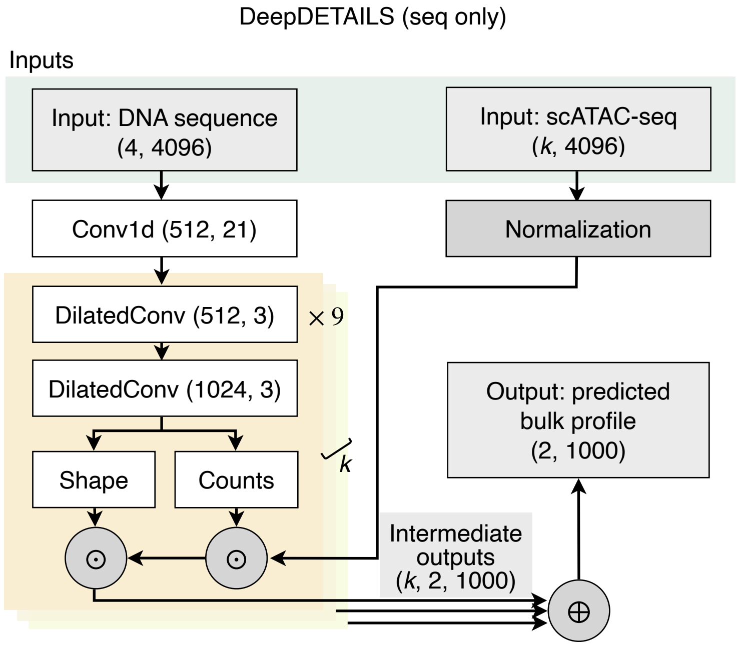


**b**

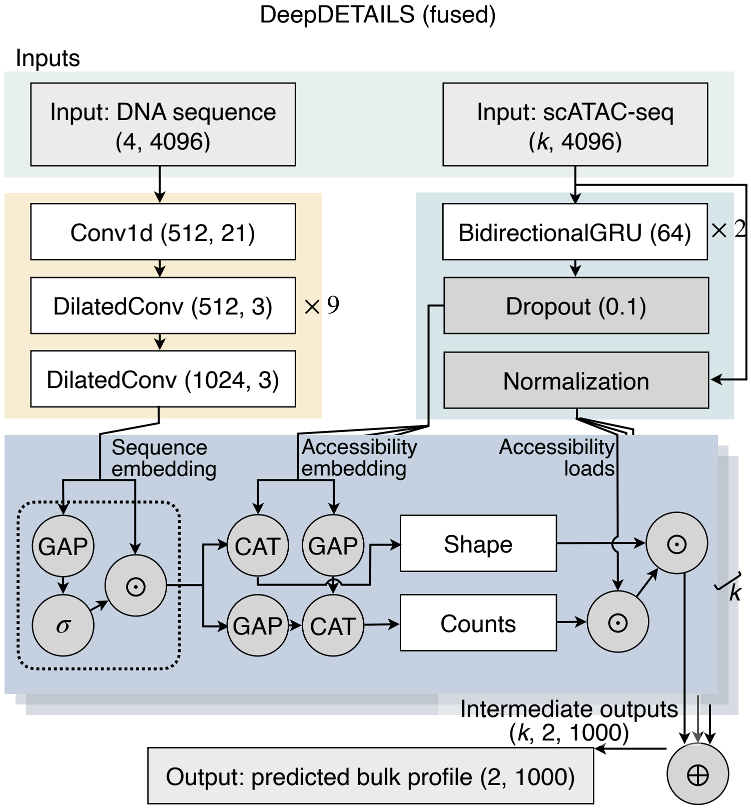

#### Supplementary Fig. 2 (continued)

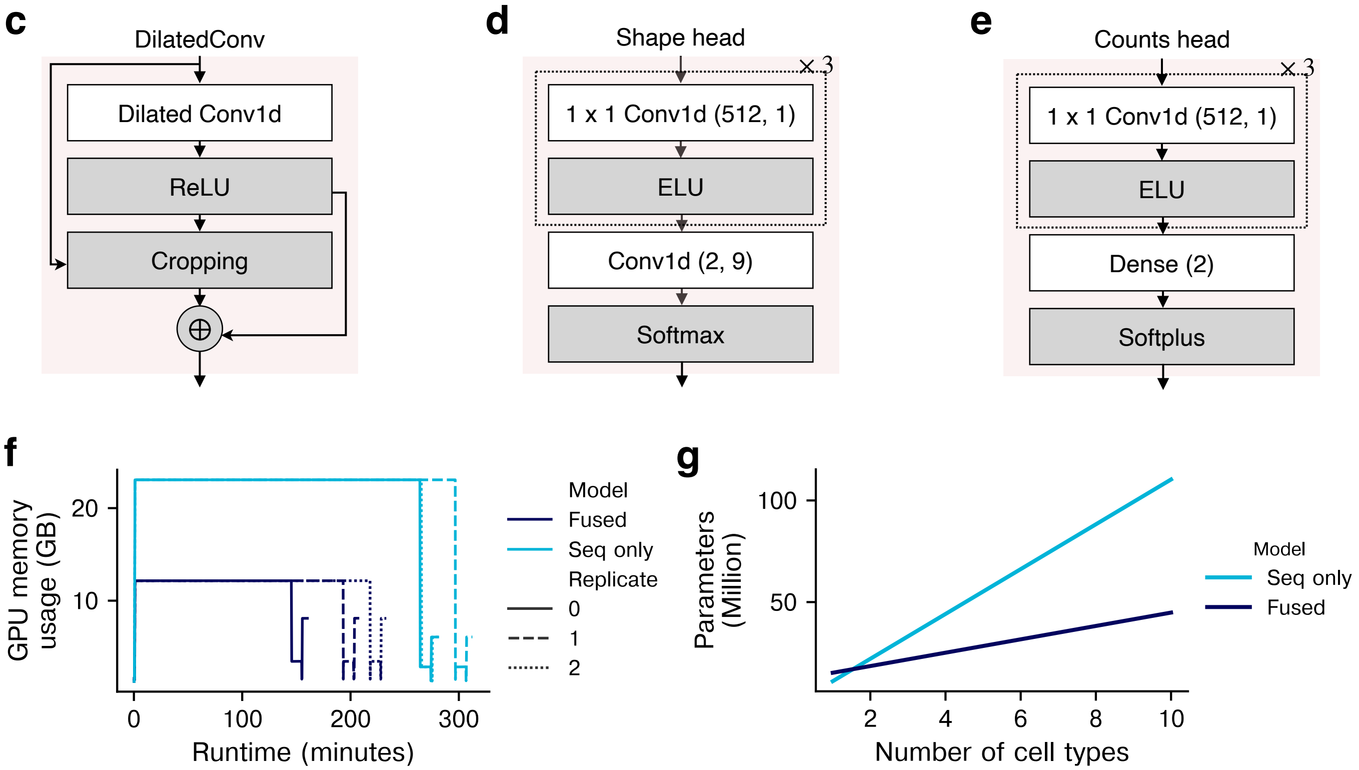

#### Supplementary Fig. 3

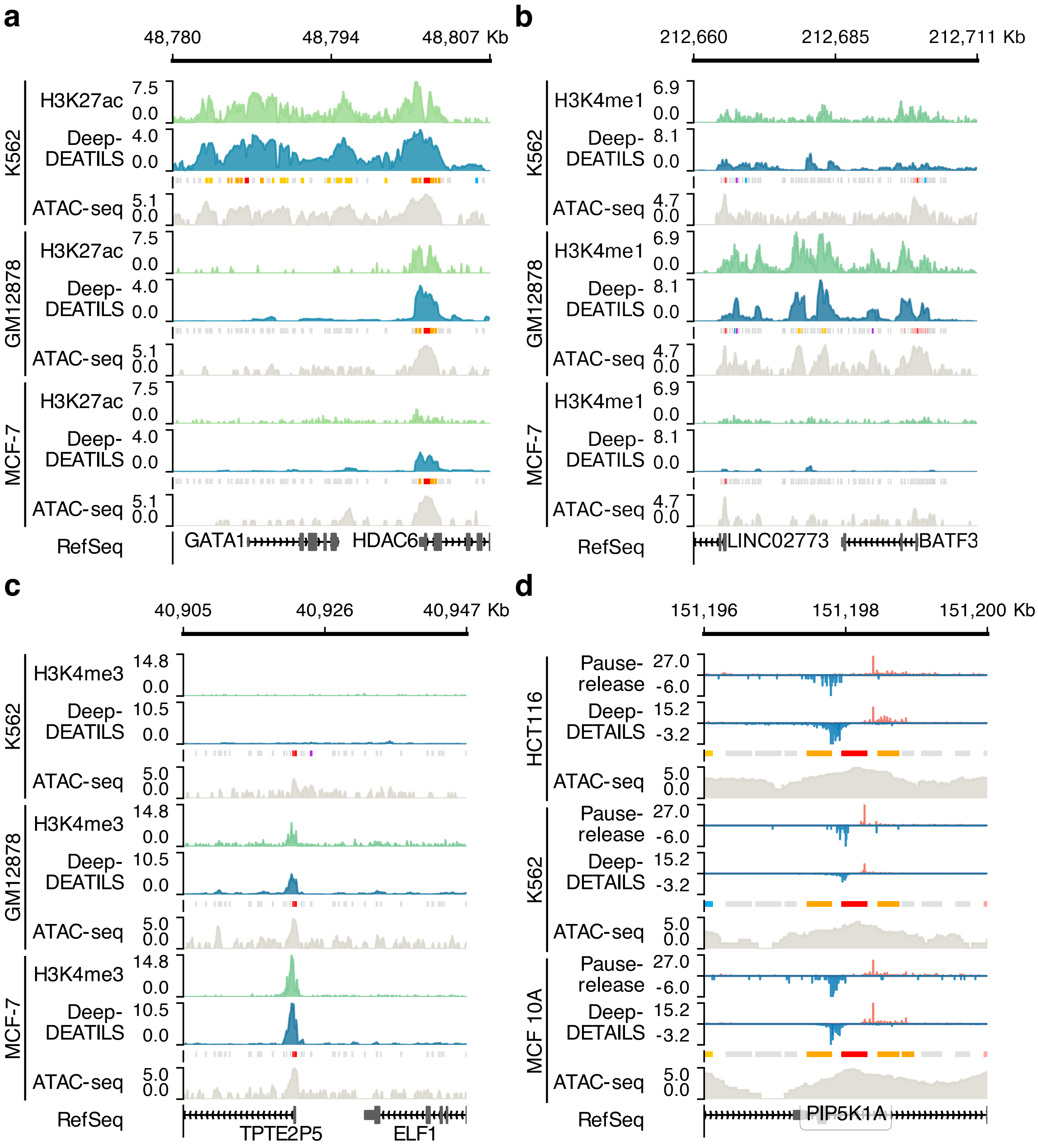

### Supplementary Fig. 4

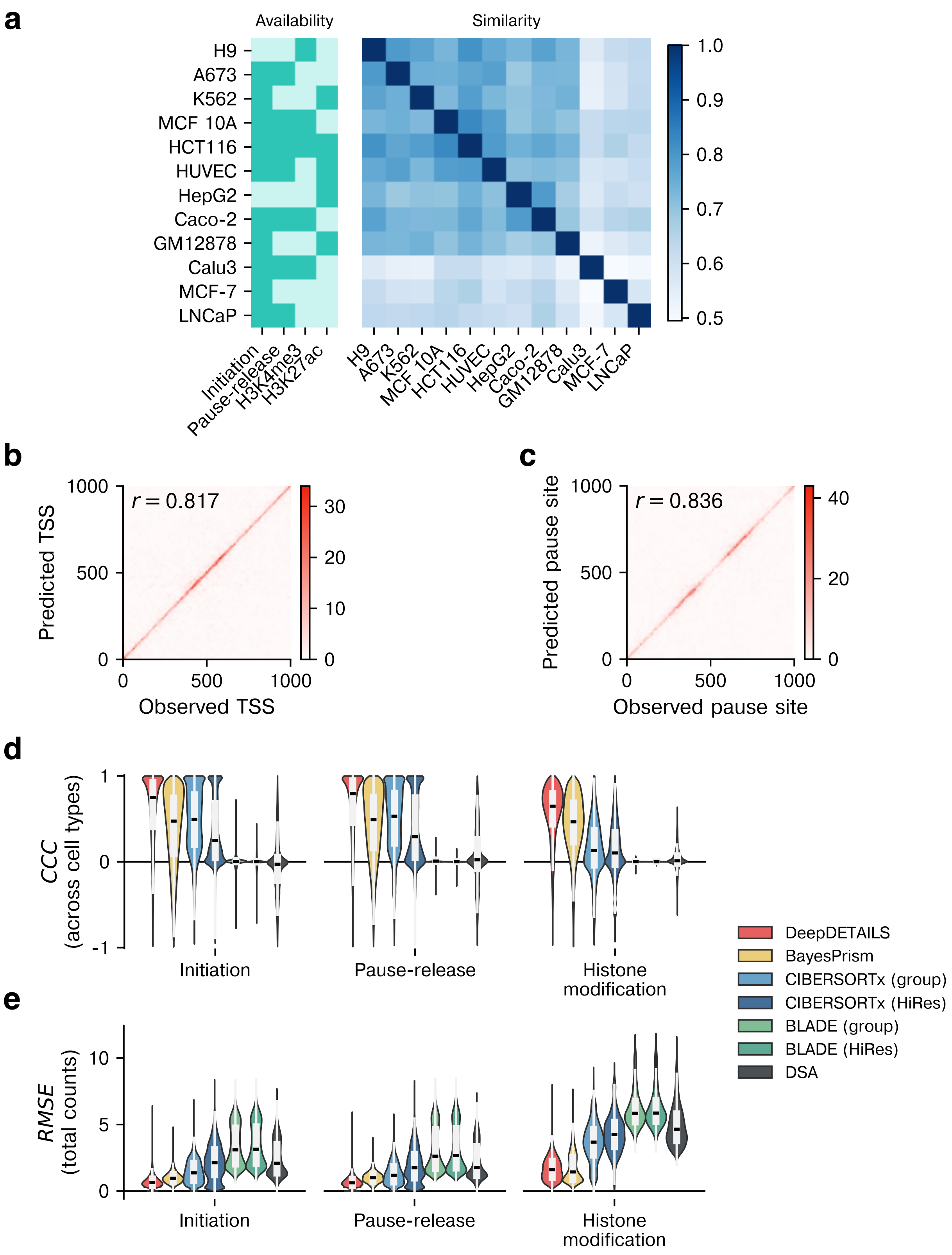

### Supplementary Fig. 5

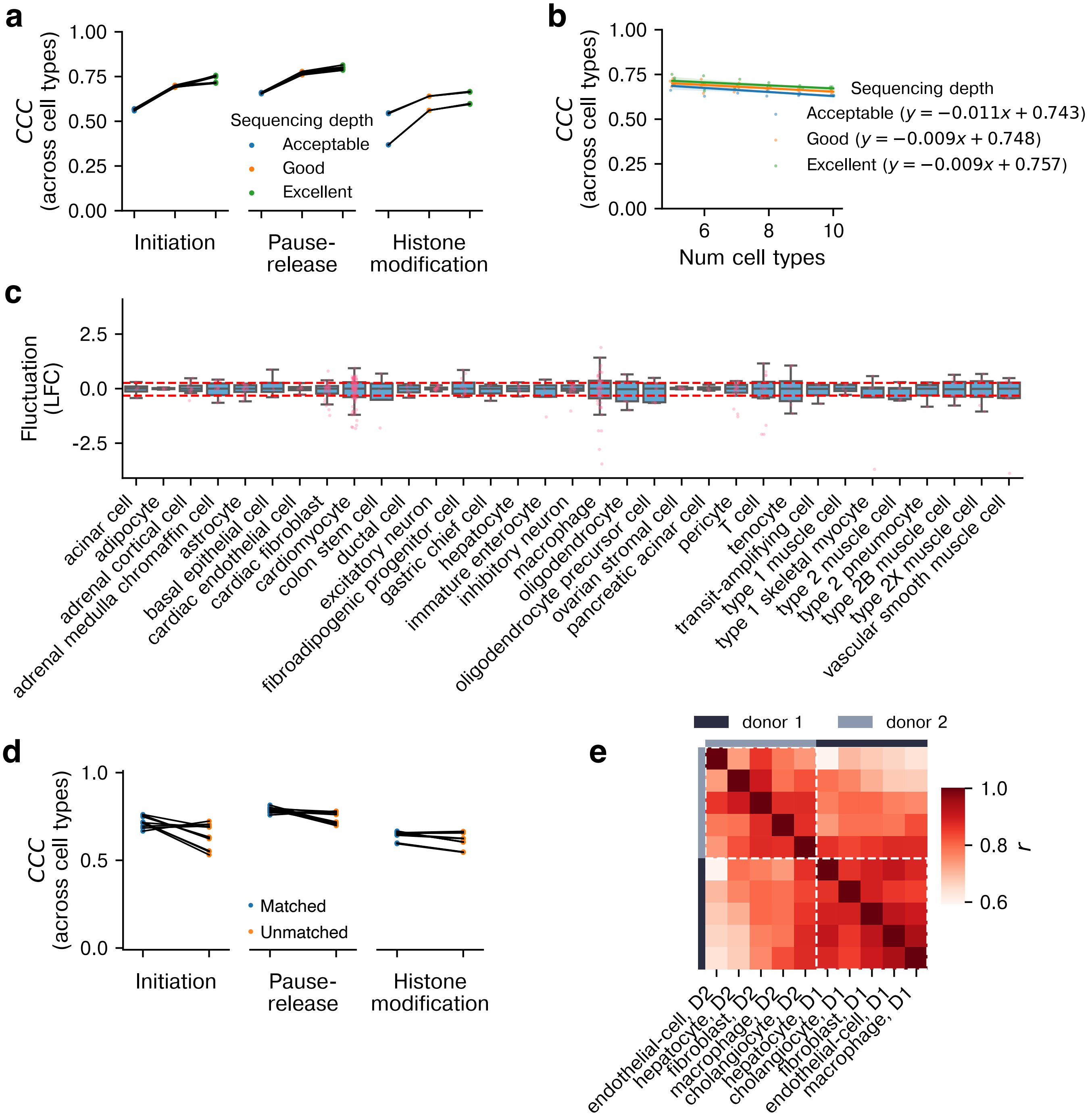

### Supplementary Fig. 6

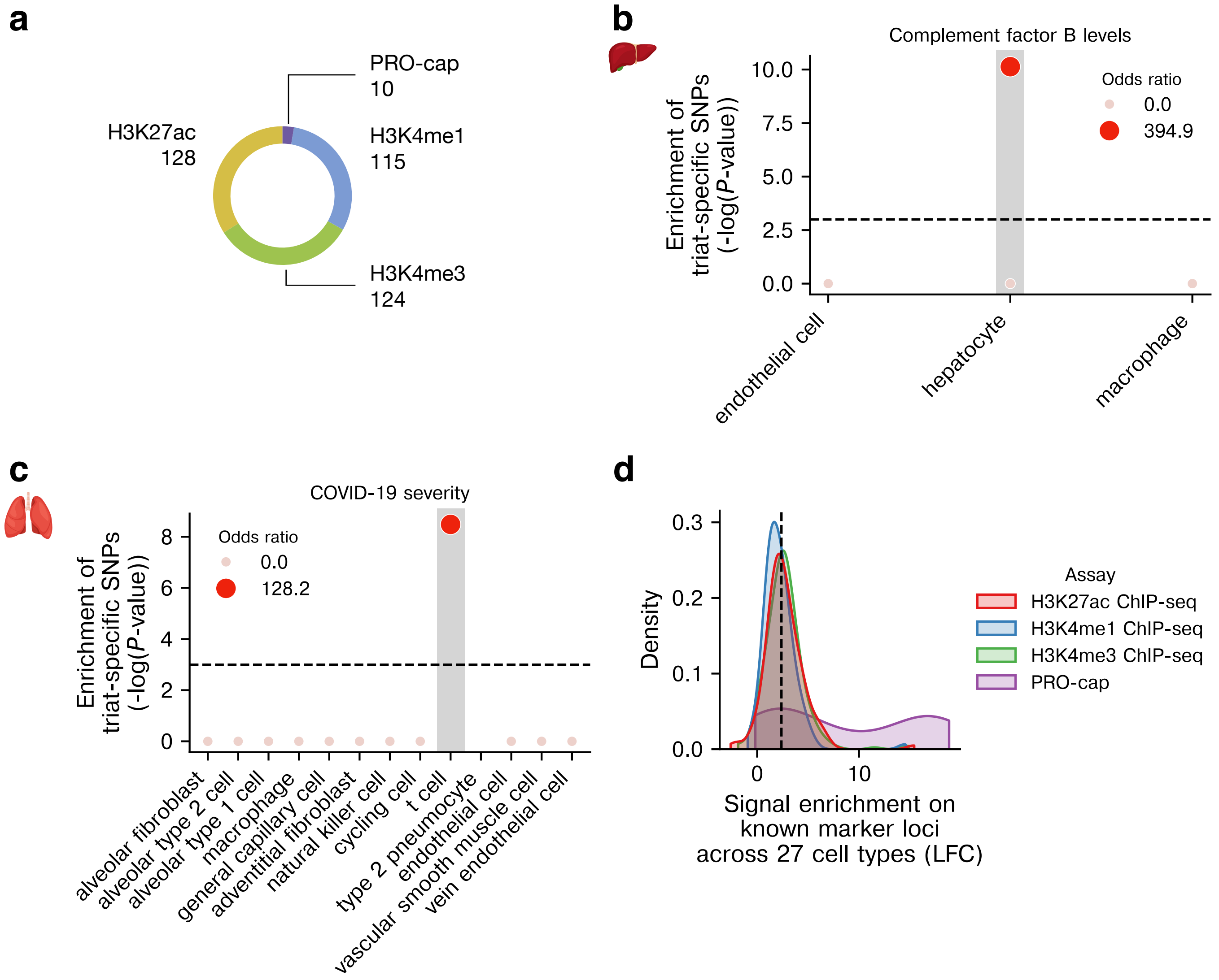

#### Supplementary Fig. 7

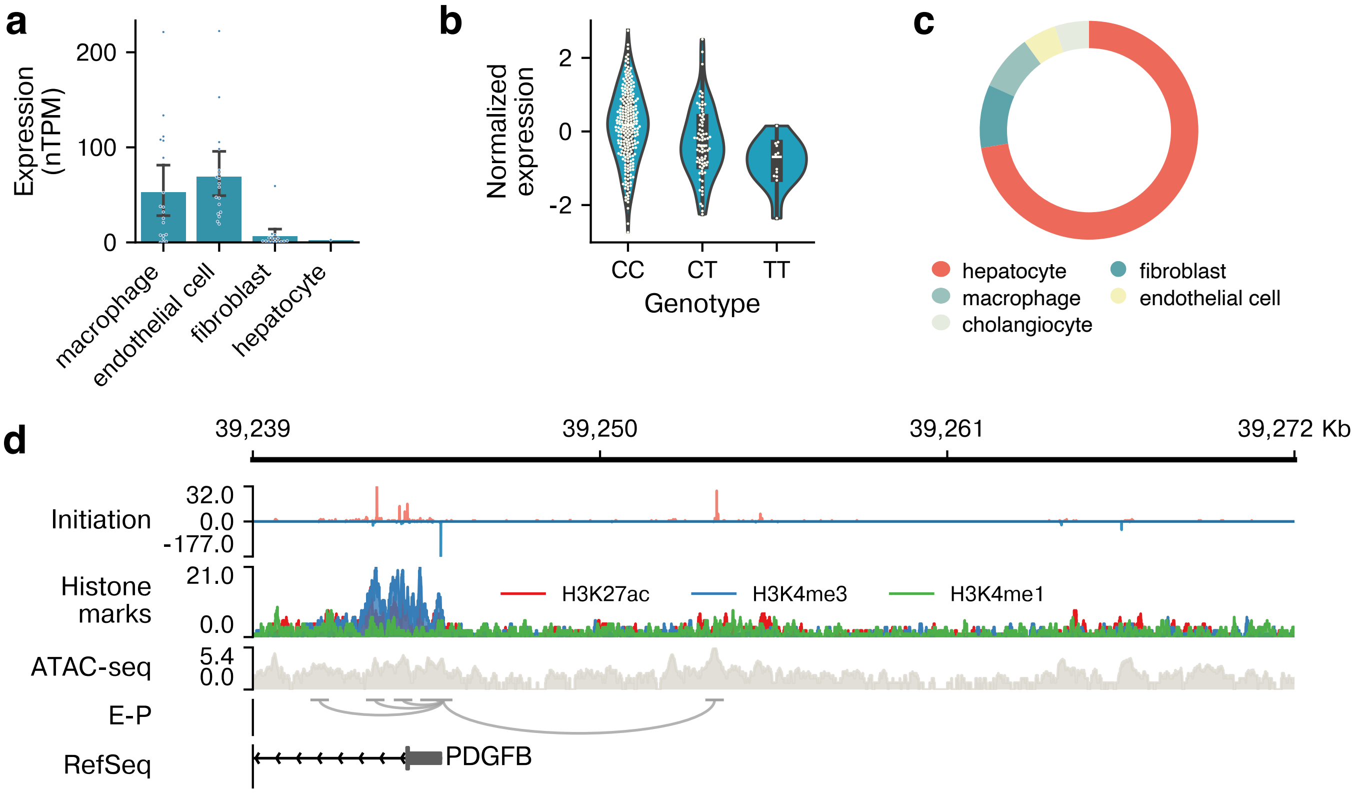
